# Complementary Organization of Driver and Modulator Cortico-Thalamo-Cortical Circuits

**DOI:** 10.1101/2024.07.14.603466

**Authors:** Rachel M. Cassidy, Angel V. Macias, Willian N. Lagos, Chiamaka Ugorji, Edward M. Callaway

## Abstract

Corticocortical (CC) projections in the visual system facilitate the hierarchical processing of sensory information. In addition to direct CC connections, indirect cortico-thalamo-cortical (CTC) pathways through the pulvinar nucleus of the thalamus can relay sensory signals and mediate interactions between areas according to behavioral demands. While the pulvinar is extensively connected to the entire visual cortex, it is unknown whether transthalamic pathways link all cortical areas or whether they follow systematic organizational rules. Because pulvinar neurons projecting to different cortical areas are spatially intermingled, their input/output relationships have been difficult to characterize using traditional anatomical methods. To determine the organization of CTC circuits, we mapped the higher visual areas (HVAs) of mice with intrinsic signal imaging and targeted five pulvinar→HVA pathways for projection-specific rabies tracing. We aligned post-mortem cortical tissue to *in vivo* maps for precise quantification of the areas and cell types projecting to each pulvinar→HVA population. Layer 5 corticothalamic (L5CT) “driver” inputs to the pulvinar originate predominantly from primary visual cortex (V1), consistent with the CC hierarchy. L5CT inputs from lateral HVAs specifically avoid driving reciprocal connections, consistent with the “no-strong-loops” hypothesis. Conversely, layer 6 corticothalamic (L6CT) “modulator” inputs are distributed across areas and are biased toward reciprocal connections. Unlike previous studies in primates, we find that every HVA receives disynaptic input from the superior colliculus. CTC circuits in the pulvinar thus depend on both target HVA and input cell type, such that driving and modulating higher-order pathways follow complementary connection rules similar to those governing first-order CT circuits.

**Significance Statement:** Understanding the functional role of the visual pulvinar will require knowledge of its anatomical connections. Using state-of-the-art rabies tracing, we establish a comprehensive map of brain-wide and CTC pulvinar connections. While the tectopulvinar pathway in primates selectively targets dorsal visual areas, we find SC input to every projection population in the mouse. This extrageniculate projection can support unconscious visually-guided behavior, suggesting that in mice all visual cortical areas contribute to such functions. Our results also unify several longstanding theories of thalamocortical anatomy. Namely, “driver” CTC inputs are feedforward relays that mirror the CC hierarchy and adhere to the “no-strong-loops” hypothesis. “Modulator” L6CT inputs to pulvinar→HVA projections are overrepresented in the target HVA, reflecting the reciprocal connectivity described in previous bulk tracing studies. Together, these findings constitute a definitive and comprehensive map which can inform future physiological experiments and theoretical models of pulvinar function.

## Introduction

An animal’s perception and interactions with the external world rely on incoming sensory information. In the mammalian visual system, the dorsal lateral geniculate nucleus (dLGN) relays this information to the neocortex, where a hierarchically organized network processes increasingly complex features. This hierarchy is defined by the laminar organization of direct CC projections (Felleman & Van Essen, 1991; Wang et al., 2012). However, the dLGN is not the only thalamocortical (TC) input. In fact, all visual areas receive projections from the pulvinar, a higher-order thalamic nucleus whose primary input is the cortex itself. This bidirectional connectivity could allow the pulvinar to route sensory information between cortical areas independently from the direct pathway (Sherman & Guillery, 2011). The pulvinar also relays ascending signals from the superior colliculus (SC) and retina (Allen et al., 2016; Beltramo & Scanziani, 2019; Lyon et al., 2010; N. Zhou et al., 2017). Without a comprehensive map of synaptic connections in the pulvinar, however, it is unclear how these transthalamic circuits relate to the hierarchy in visual cortex (Halassa & Sherman, 2019).

Available viral and genetic tools make the mouse an ideal model for dissecting fine-scale input/output connections. Murine visual cortex includes V1 and multiple higher visual areas (HVAs) with hierarchically organized feedforward and feedback connections (Garrett et al., 2014; Harris et al., 2019; Wang et al., 2012; Wang & Burkhalter, 2007). In rodents, individual pulvinar (or lateral posterior nucleus) neurons project to the input layers of only 2-3 cortical areas (Nakamura et al., 2015). While pulvinar outputs are not completely independent, then, they do not broadcast indiscriminately (Blot et al., 2021; Juavinett et al., 2020). Therefore, TC projections could send distinct signals to HVAs depending on the organization of their corticothalamic (CT) inputs. CT axons are organized topographically in the pulvinar, with distributions that roughly align with the cell bodies of corresponding TC projections (Bennett et al., 2019; Juavinett et al., 2020; Tohmi et al., 2014). While this coarse input/output topography is reciprocal, regions of overlap could still support interareal relay (Sherman & Guillery, 2011; Shipp, 2003). For a transthalamic pathway to transmit sensory information, however, CT neurons must be capable of driving activity. In the pulvinar, layer 5 (L5CT) “drivers” are necessary for visual responses and receptive field structure, whereas layer 6 (L6CT) “modulators” are not required for visual activity, but can affect the frequency and gain of responses (Kirchgessner et al., 2021; Sherman, 2016). Therefore, describing both the cortical regions and cell types connecting to each pulvinar→HVA population is prerequisite to understanding cortico-thalamo-cortical (CTC) circuits.

Determining fine-scale CTC circuit organization requires projection-specific tracing of the relationships between pulvinar inputs and outputs (“TRIO”; Schwarz et al., 2015). Previous studies applied this technique to a limited number of pulvino-cortical projections (Blot et al., 2021; Leow et al., 2022; Miller-Hansen & Sherman, 2022), but it is unclear whether those findings can be generalized. To reveal the organizational principles governing CTC connections across the visual hierarchy, we targeted pulvinar axons in each of five functionally mapped HVAs (LM, AL, RL, AM, and PM) for Cre-dependent, monosynaptic rabies tracing and quantified their brain-wide inputs. Consistent with previous results of AL and PM projections, we find that V1 provides the largest source of driving L5CT input to every pulvinar→HVA pathway, confirming the long-standing theory that CTC connections serve as a parallel feedforward relay (Blot et al., 2021; Sherman & Guillery, 2011). L5CT pathways from HVAs specifically avoid reciprocal connections, consistent with the “no-strong-loops” hypothesis (Crick & Koch, 1998). In contrast, L6CT inputs are distributed across V1 and HVAs, and they preferentially form reciprocal CTC loops. Driving and modulating inputs to the pulvinar thus form complementary anti-reciprocal and reciprocally-biased pathways, respectively. Regardless of the target HVA, all pulvinar projections receive SC input. These results show that the pulvinar is organized as a feedforward pathway between cortical areas that reflects the directionality of the existing cortical hierarchy, and that each of these parallel pathways could integrate sensorimotor information from the SC.

## Results

### Brain-wide mapping of pulvinar input/output connections

We characterized the brain-wide inputs to specific pulvinar→HVA pathways using projection-specific, Cre-dependent, monosynaptic rabies tracing (Fig. 1). After Cre-dependent AAV helper viruses (AAV8-Flex-TVAmCherry and AAV8-DIO-oG) were injected into the pulvinar of wild-type mice (n = 24, 4-5 per target HVA; See Table 1), we used intrinsic signal imaging (ISI) to generate retinotopically defined borders of five HVAs (PM, AM, RL, AL, and LM) (Fig. 1a,b). These functional maps guided precise, restricted injections of AAV5-Cre into each HVA, which retrogradely infected pulvinar neurons and drove Cre-dependent expression of the avian TVA receptor and rabies glycoprotein (oG) (Fig. 1b). EnvA+RVΔG-eGFP was injected into the same location in pulvinar after 21 days. Animals were sacrificed 10 days later to allow for sufficient transsynaptic rabies labeling. Because HVA borders vary between individual animals, we aligned each functionally defined ISI map to post-mortem, tangentially-sectioned cortical tissue for quantification of GFP-expressing inputs (Fig. 1b,d). Non-cortical brain areas were also sectioned and processed for quantification of brain-wide inputs (Fig. 1e). Cre-dependent rabies tracing in reciprocally connected brain areas requires careful selection of viral serotypes and concentrations to avoid both false positive and false negative labeling of input cells (Lavin et al., 2020).

**Figure 1.**
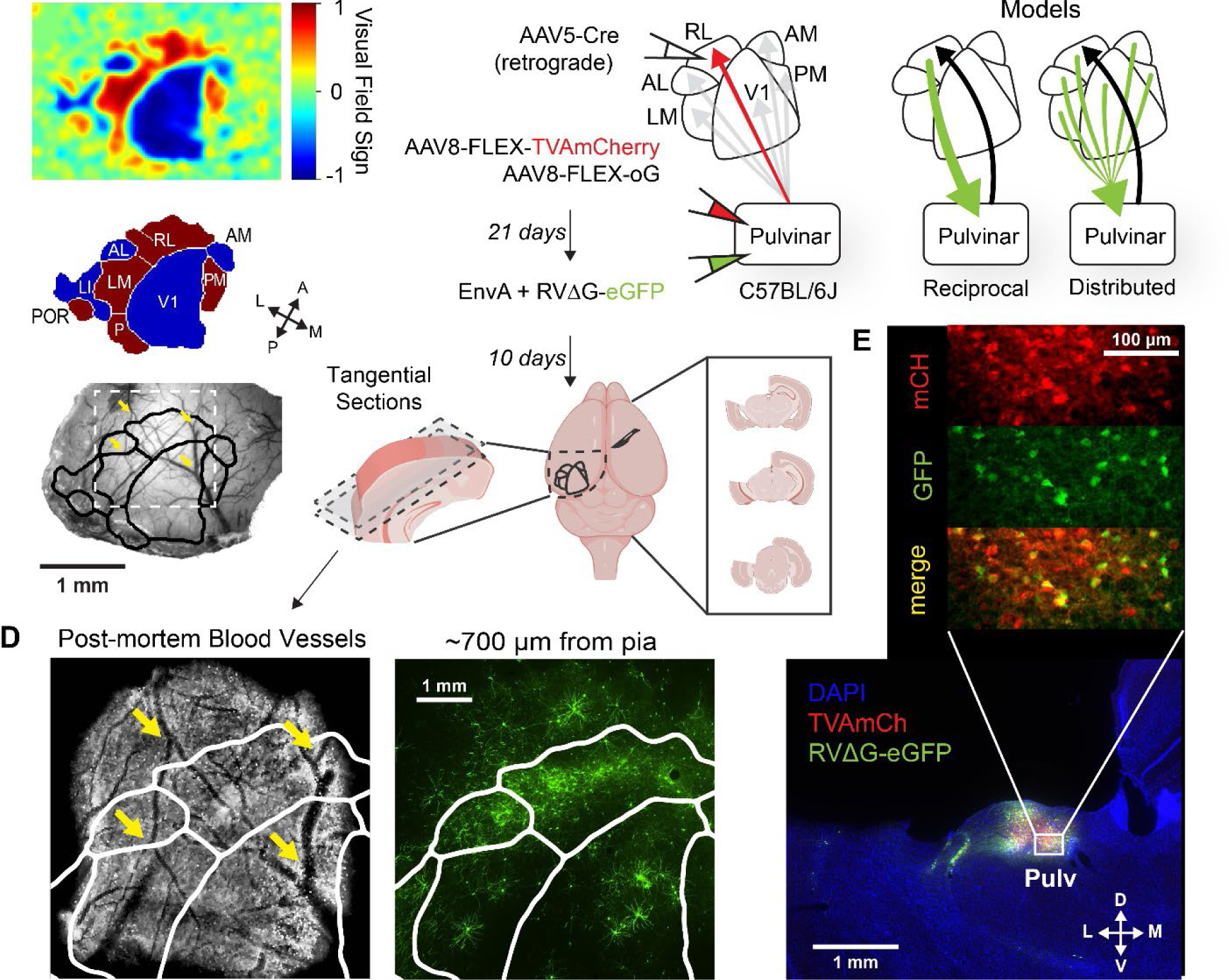
Targeting visual areas for pulvino-cortical input tracing. A) Intrinsic signal imaging and automatic segmentation defines HVA borders aligned to *in vivo* blood vessels (animal RL2). B) Schematic of viral injection strategy for projection-specific input tracing. (top) AAV5-Cre injected into individual HVAs in wild-type mice retrogradely infects pulvinar projection neurons expressing Cre-dependent helper viruses. Pseudotyped, G-deleted rabies expressing eGFP is injected into the pulvinar to label presynaptic inputs to specific pulvino-cortical projection populations. (bottom) Visual cortex is dissected and sectioned tangentially to align tissue with the *in vivo* blood vessels and HVA map. The remainder of the tissue is sectioned coronally and processed to quantify brain-wide inputs. C) Models of cortico-thalamo-cortical connectivity. D) Tangential cortical sections aligned to the HVA map. (left) Surface blood vessel landmarks in post-mortem tissue are aligned with the in vivo imaging. (right) GFP+ cortico-pulvinar inputs in aligned deeper sections. E) Putative starter cells in the pulvinar expressing TVAmCherry and RVΔG-eGFP.

**Table 1.** Summary of Input and Starter Cell Counts.

| Target HVA | Animal ID | Total Input Neurons | Number of Starter Neurons by Thalamic Nucleus |  |  |  |  |
| --- | --- | --- | --- | --- | --- | --- | --- |
|  |  |  | Pulvinar/LP | LD | POm | dLGN | CL |
| PM | PM1 | 7710 | 141<br>(92.2%) | 4<br>(2.6%) | 2<br>(1.3%) | 2<br>(1.3%) | 4<br>(2.6%) |
|  | PM2 | 4562 | 403<br>(85.2%) | 45<br>(9.5%) | 1<br>(0.2%) | 20<br>(4.2%) | 1<br>(0.2%) |
|  | PM3 | 2870 | 835<br>(85.2%) | 122<br>(12.5%) | 0 (0%) | 10<br>(1.0%) | 7<br>(0.1%) |
|  | PM4 | 1163 | 666<br>(73.4%) | 224<br>(24.7%) | 5<br>(0.6%) | 2<br>(0.2%) | 8<br>(0.9%) |
|  | PM5 | 4505 | 377<br>(73.8%) | 74<br>(14.5%) | 19<br>(3.7%) | 5<br>(1.0%) | 26<br>(5.1%) |
| AM | AM1 | 1531 | 313<br>(59.3%) | 203<br>(38.4%) | 0 (0%) | 11<br>(2.1%) | 0 (0%) |
|  | AM2 | 5182 | 337<br>(87.1%) | 32<br>(8.3%) | 2<br>(0.5%) | 6<br>(1.6%) | 8<br>(2.1%) |
|  | AM3 | 1707 | 292<br>(92.4%) | 5<br>(1.6%) | 17<br>(5.4%) | 2<br>(0.6%) | 0 (0%) |
|  | AM4 | 2585 | 158<br>(55.4%) | 44<br>(15.4%) | 58<br>(20.4%) | 4<br>(1.4%) | 14<br>(4.9%) |
|  | AM5 | 2053 | 200<br>(86.6%) | 15<br>(6.5%) | 10<br>(4.3%) | 4<br>(1.7%) | 0 (0%) |
| RL | RL1 | 5762 | 527<br>(74.3%) | 150<br>(21.2%) | 12<br>(1.7%) | 14<br>(2.0%) | 6<br>(0.8%) |
|  | RL2 | 9173 | 1058<br>(86.0%) | 148<br>(12.0%) | 19<br>(1.5%) | 2<br>(0.2%) | 4<br>(0.3%) |
|  | RL3 | 972 | 300<br>(86.2%) | 40<br>(11.5%) | 0 (0%) | 5<br>(1.4%) | 2<br>(0.6%) |
|  | RL4 | 2401 | 339<br>(73.9%) | 70<br>(15.3%) | 39<br>(8.5%) | 3<br>(0.7%) | 5<br>(1.1%) |
| AL | AL1 | 8874 | 1315<br>(91.1%) | 82<br>(5.7%) | 6<br>(0.4%) | 27<br>(1.9%) | 8<br>(0.6%) |
|  | AL2 | 2957 | 819<br>(92.3%) | 24<br>(2.7%) | 1<br>(0.1%) | 1<br>(0.1%) | 41<br>(4.6%) |
|  | AL3 | 3273 | 678<br>(83.3%) | 36<br>(4.4%) | 39<br>(4.8%) | 52<br>(6.4%) | 1<br>(0.1%) |
|  | AL4 | 2187 | 155<br>(68.3%) | 5<br>(2.2%) | 56<br>(24.7%) | 3<br>(1.3%) | 0 (0%) |
|  | AL5 | 4258 | 368<br>(92.7%) | 0 (0%) | 16<br>(4.0%) | 8<br>(2.0%) | 0 (0%) |
| LM | LM1 | 2595 | 117<br>(95.9%) | 0 (0%) | 0 (0%) | 3<br>(2.5%) | 1<br>(0.8%) |
|  | LM2 | 1670 | 85 (100%) | 0 (0%) | 0 (0%) | 0 (0%) | 0 (0%) |
|  | LM3 | 2373 | 88 (75.2%) | 10<br>(8.5%) | 2<br>(1.7%) | 9<br>(7.7%) | 4<br>(3.4%) |
|  | LM4 | 2444 | 544<br>(81.1%) | 56<br>(8.3%) | 12<br>(1.8%) | 28<br>(4.2%) | 30<br>(4.5%) |
|  | LM5 | 3069 | 638<br>(77.9%) | 100<br>(12.2%) | 0 (0%) | 78<br>(9.5%) | 1<br>(0.1%) |

To optimize combinations of AAVs, we conducted numerous troubleshooting and control experiments (See Methods and Fig. 2). In our hands, cortical injections of AAVretro-Cre, commonly used for retrograde targeting (Tervo et al., 2016), resulted in pronounced tissue damage and complete absence of rabies-labeled CT neurons at the injection site. Lowering the concentration of AAVretro-Crereduced cortical damage, but Cre expression in the pulvinar with sufficiently diluted AAVretro was equivalent to or lower than with injections of AAV5-Cre. To rule out Cre-independent rabies spread, we injected helper AAVs and EnvA+RVΔG-eGFP while omitting AAV5-Cre and confirmed that no cells outside of the immediate injection site expressed GFP. In reciprocally connected thalamocortical circuits, however, AAV-FLEx-TVA injections into the pulvinar can mediate retrograde infection of Cre-expressing CT neurons in the target area, which are then capable of expressing TVA at levels permitting direct uptake of EnvA+RVΔG from axon terminals in the pulvinar. Because this glycoprotein-independent labeling is facilitated by Cre expression, it is essential to conduct control experiments *with* AAV-Cre while omitting Cre- dependent AAV-oG.

**Figure 2.**
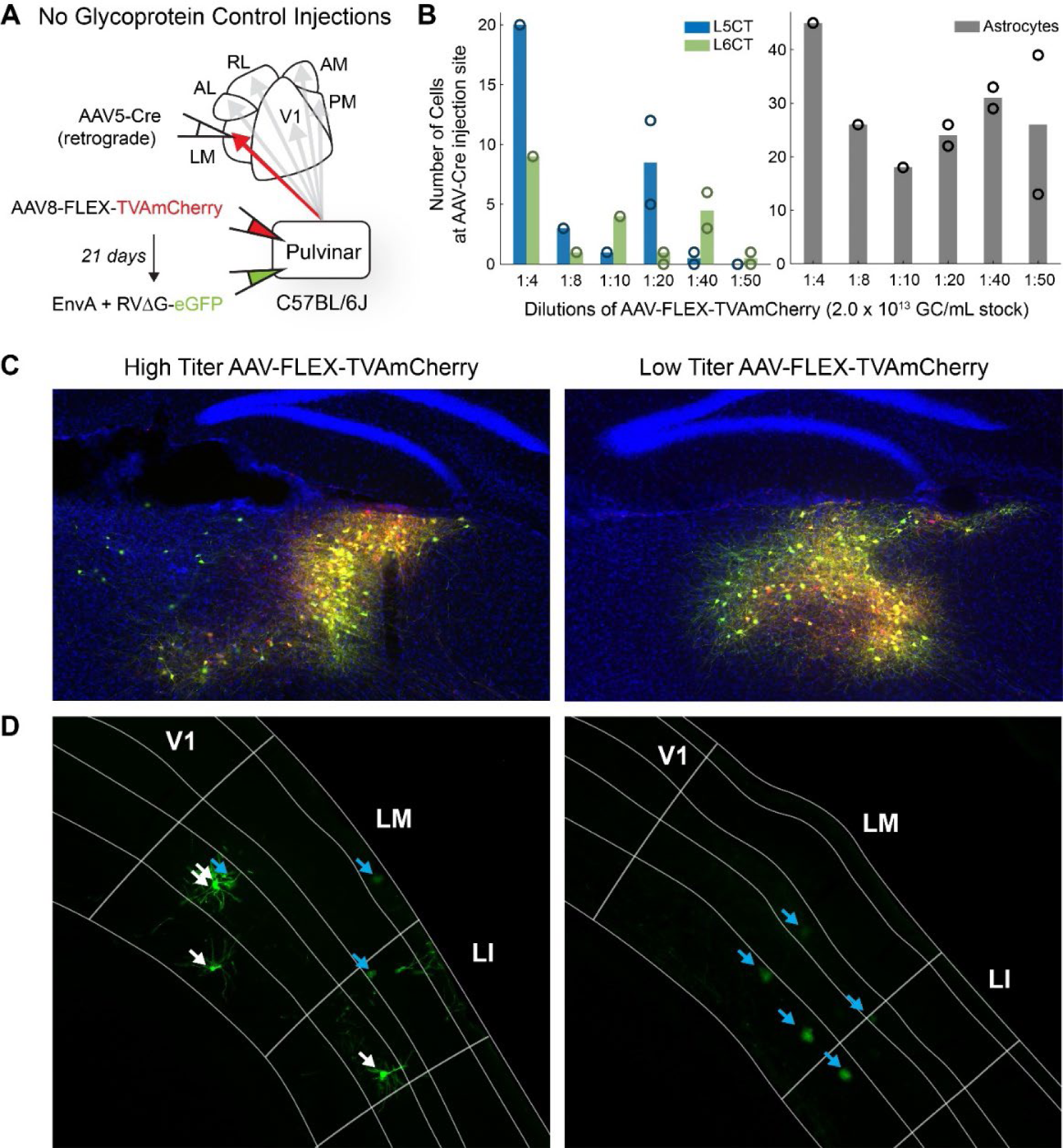
High titer AAV-FLEx-TVA permits retrograde, glycoprotein-independent rabies spread to CT neurons expressing Cre. A) Schematic of viral injection strategy for “no glycoprotein” control injections. **B)** Number of GFP-labeled CT neurons (left) and astrocytes (right) at the cortical AAV-Cre injection site with varying AAV-FLEx-TVAmCherry titer. Bars indicate the mean cell counts. Open circles represent counts for individual brains (1:4-1:10 dilutions: n = 1, 1:20-1:50 dilutions: n = 2) **C)** LM-projecting pulvinar neurons express TVAmCherry (red) and RVdG-eGFP (green) for high (5.1 x 10^12^ GC/mL; left) and low (4.1 x 10^11^ GC/mL; left) titer AAV-FLEx-TVAmCherry injections. **D)** In the absence of TC glycoprotein expression, LM astrocytes (blue arrows) express GFP for both high (left) and low (right) titer AAV-FLEx-TVAmCherry injections, but CT pyramidal neurons (white arrows) express GFP only when AAV-FLEx-TVAmCherry titer is high (left).

In initial tests, we frequently observed GFP+ CT neurons concentrated at the AAV-Cre injection site. These cells almost never had a detectable mCherry signal, underscoring that the lack of visible TVA expression is an inadequate quality control criterion. In a comparison of different helper virus serotypes, we found that AAV8 injections in the pulvinar mediated the least amount of retrograde uptake from CT neurons. Using two separate AAVs for TVA and oG rather than a bicistronic vector, we titrated the concentration of AAV8-FLEx-TVAmCherry until control injections without AAV8-DIO-oG resulted in negligible RV labeling in the target cortical area (Fig. 2).

Using the reagents and viral titers established in control experiments, we observed putative starter cells in the pulvinar consistent with the previously reported topography of HVA projections (Bennett et al., 2019; Juavinett et al., 2020). The starter cell coverage was sensitive to the volume and stereotaxic coordinates of helper virus injections, such that small or off target injections led to biased labeling of starter cells.

Total input neurons for individual animals are shown. Starter neuron counts in the top five thalamic nuclei alongside the corresponding percentage of total starter neurons. LP: lateral posterior nucleus; LD: lateral dorsal nucleus; POm: posteromedial nucleus; dLGN: dorsal lateral geniculate nucleus; CL: central lateral nucleus.

To ensure that each HVA projection was adequately sampled, we injected a larger volume of helper virus that spread beyond the borders of the pulvinar (See Methods). We excluded off-target injections with incomplete coverage of the pulvinar compared to unbiased distributions (Bennett et al., 2019).Because the proportion of starter cells in the pulvinar was 82.0 ± 2.3% (mean ± SEM, Table 1), we consider the resulting inputs to largely reflect the connections of the pulvinar nucleus. However, a small proportion of starter cells were located in adjacent higher-order thalamic nuclei such as the laterodorsal (LD) and the posteromedial (POm) nuclei, which also project to HVAs.

Figure 3 illustrates the input from all major brain regions to pulvinar neurons projecting to each of 5 HVAs expressed as a proportion of all rabies labeled input neurons. Input from each of these areas did not depend on the cortical target (p>0.05), except for SC (p = 6.9 x 10^-3^, Kruskal-Wallis test). Consistent with previous bulk retrograde tracing, we found that the largest source of input to the pulvinar pooled across projection groups was the cortex (41.4 ± 1.3% mean ± SEM), followed by the thalamic reticular nucleus (15.1 ± 0.9% mean ± SEM), SC (11.0 ± 1.3% mean ± SEM) and pretectal nuclei (10.8 ± 0.6% mean ± SEM). and cortical inputs.

**Figure 3.**
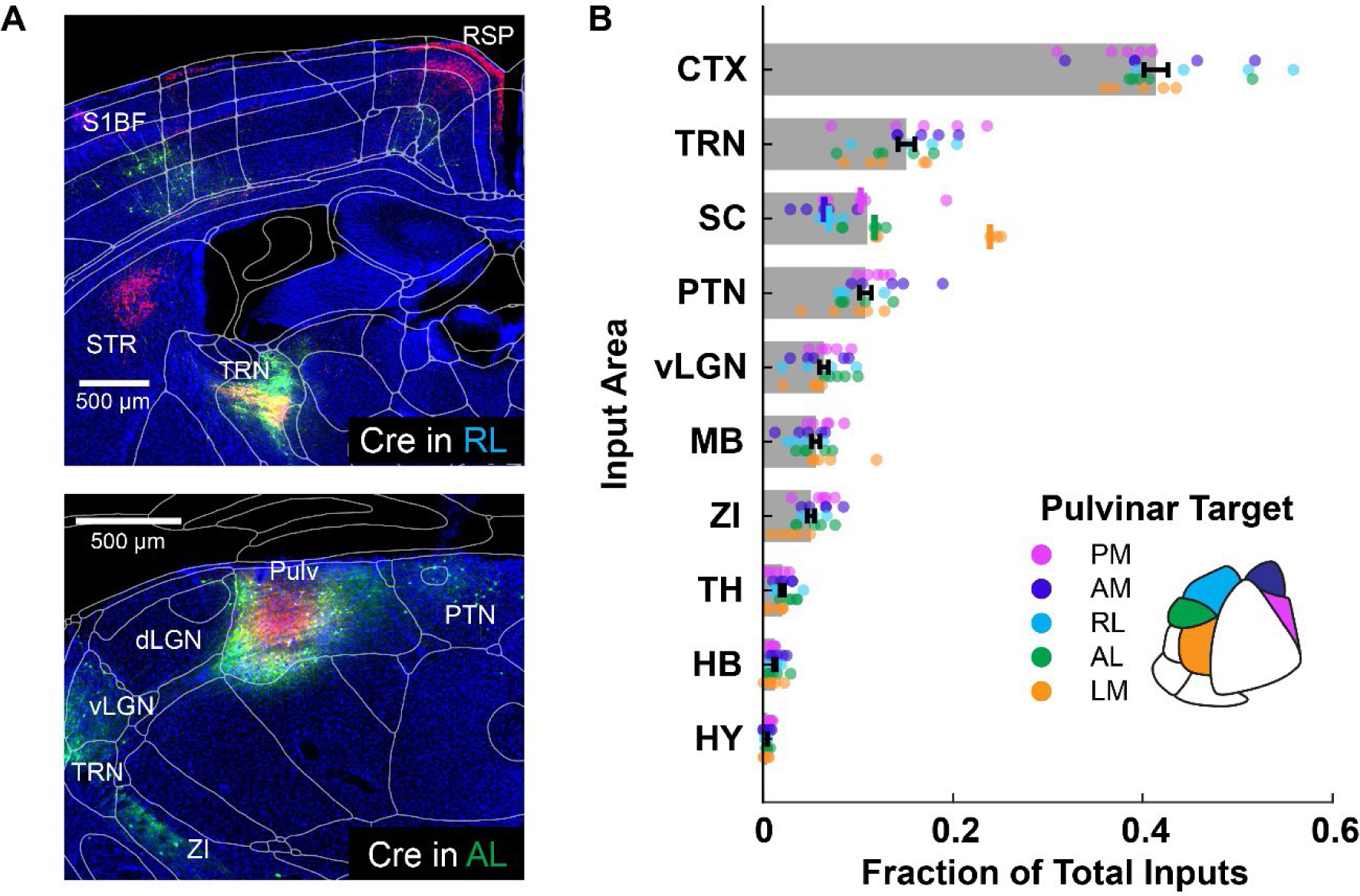
Brain-wide inputs to the pulvinar are similar for different projection targets. A) Example coronal sections of Cre-dependent rabies labeling in two example experiments with transformed CCF atlas borders. Scale bars = 500 μm. *Top*: RVΔG-eGFP+ inputs (green) in CTX and TRN and TVAmCherry+ pulvinar axons (red) in CTX, STR, and TRN are labeled in animal RL2. *Bottom*: Starter cells projecting to AL are labeled in the pulvinar with TVAmCherry (red) and RVΔG-eGFP (green) in animal AL1. **B)** Brain-wide inputs to the pulvinar from major divisions of the brain, expressed as a fraction of the total inputs. Markers represent individual animals, colored according to the pulvinar projection target. Bar lengths represent the mean of the sample obtained by pooling projection targets together. For all areas except for SC, the distribution of inputs for different projection targets was not significantly different (p>0.05, kruskal-wallis test), and samples were combined for a pooled mean ± SEM represented by black error bars. For areas SC, the median input for each projection target is displayed by a vertical line. CTX: Cortex; dLGN: dorsal subdivision of the lateral geniculate nucleus; HB: hindbrain; HY: hypothalamus (other); MB: midbrain (other); PTN: pretectal nuclei; Pulv: Pulvinar; RSP: retrosplenial cortex; S1BF: primary somatosensory barrel field; SC: superior colliculus; STR: striatum; TH: thalamus (other); TRN: thalamic reticular nucleus; vLGN: ventral subdivision of the lateral geniculate nucleus; ZI: zona incerta

### The superior colliculus provides input to all projection populations

After the cortex, the SC is the second largest source of excitatory input to the pulvinar (Fig. 3b). With direct retinal input to its superficial layers, the SC forms an ascending visual pathway through the pulvinar that bypasses the dLGN and is conserved across mammalian species (Abramson & Chalupa, 1988; Harting et al., 1973; Petry & Bickford, 2019; N. Zhou et al., 2017). In macaques, this pathway projects to dorsal stream visual cortical areas (e.g. V3 and MT), but never ventral areas V2 and V4 (Lyon et al., 2010). Surprisingly, we find SC inputs to every pulvinar→HVA population in the mouse (proportion of total input from SC to pulvinar→LM: 23.9 ± 1.1%, AL: 11.7 ± 1.2%, RL: 6.9 ± 0.7%, AM: 6.4 ± 1.8%, PM: 10.3 ± 3.4% median ± median absolute deviations; Fig. 4a,b). LM-projecting pulvinar neurons receive a significantly larger proportion of their input from SC than do RL- and AM- projecting pulvinar neurons (p(LM vs. RL) = 0.04, p(LM vs. AM) = 0.006, p > 0.05 for all other pairwise comparisons, Kruskal-Wallis test with Dunn-Šidák post hoc correction). These results are consistent with previous work showing that SC targets the caudal region of the pulvinar, which in turn projects to lateral extrastriate areas (P, POR, LM, and LI) (Beltramo & Scanziani, 2019; Bennett et al., 2019; N. Zhou et al., 2017, 2018). As this pathway is topographically restricted, the SC→pulvinar→LM input in this study is likely underestimated, as thalamic injections did not always spread to the extreme caudal end of the pulvinar.

**Figure 4.**
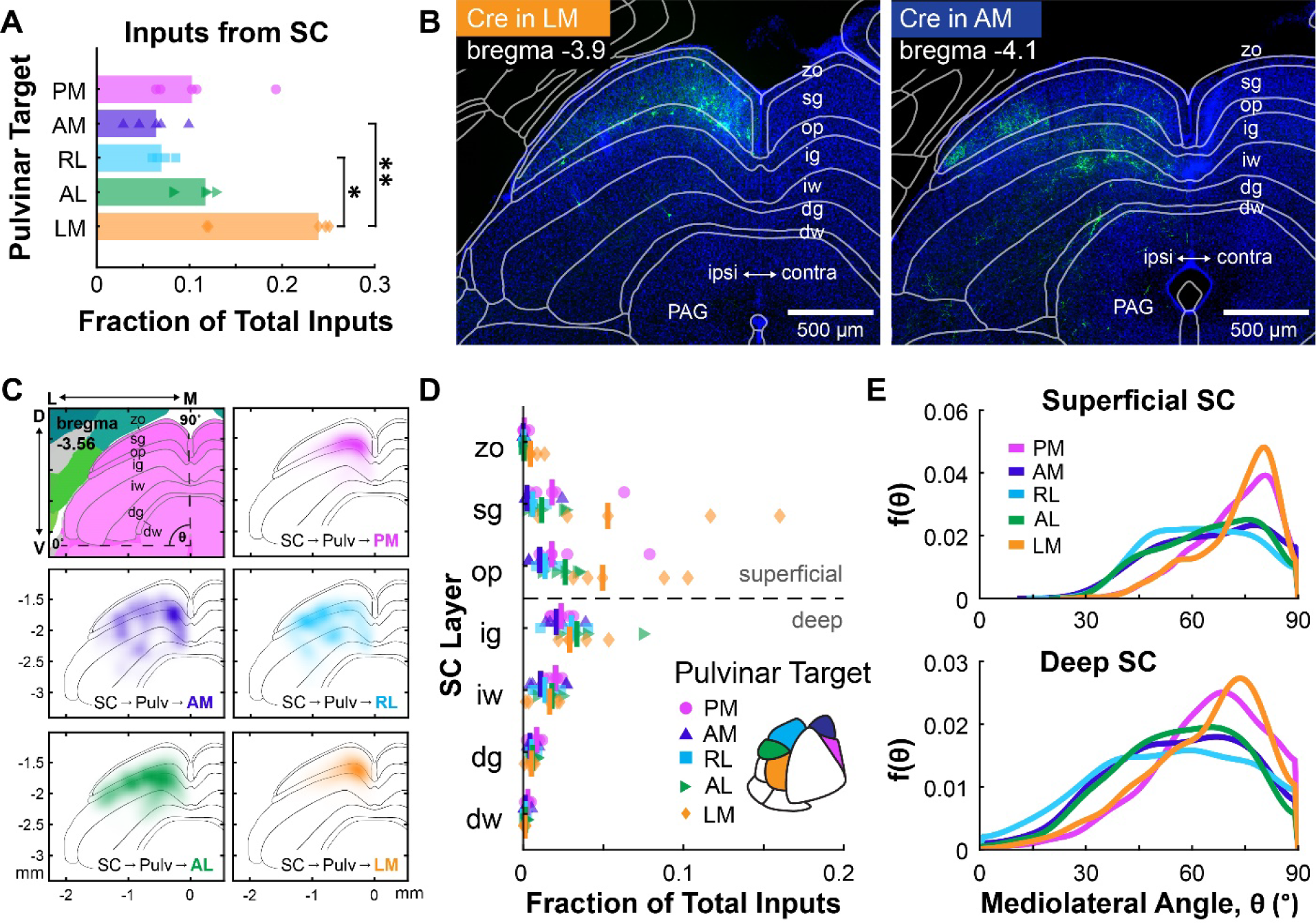
Superior Colliculus targets all pulvino-cortical projection populations. A) SC input to PM- (magenta), AM- (purple), RL- (cyan), AL- (green), and LM-projecting (orange) pulvinar neurons. Bar lengths correspond to the median SC input expressed as a fraction of total inputs. Input to LM-projecting neurons was significantly greater than to AM- (padj=0.0067) and RL- (padj=0.0434) projecting neurons (Kruskal-Wallis nonparametric rank test with Dunn-Šidák post- hoc correction for multiple comparisons). All other pairwise comparisons were nonsignificant. **B)** Example coronal sections of GFP+ inputs in the SC from experiments targeting pulvinar projections to LM (left, LM1) and AM (right, AM5). Transformed CCF borders are superimposed. Scale bars = 500 μm. zo: zonal; sg: superficial gray; op: optic; ig: intermediate gray; iw: intermediate white; dg: deep gray; dw: deep white; PAG: Periaqueductal gray. **C)** Top left: Coronal atlas of SC layers at -3.56 mm from bregma, adapted from the Allen Mouse Brain Atlas (Wang et al., 2020). Polar coordinates define mediolateral position in the SC (Benavidez et al., 2021), where θ = 0° represents the ventral border of the SC and θ = 90° represents the midline. Top right: Smoothed distribution of inputs labeled in the SC from all PM-projecting experiments combined. Middle: AM (left) and RL (right) input distributions. Bottom: AL (left) and LM (right) input distributions. **D)** SC input from each layer, expressed as a fraction of total inputs. Each point at each depth corresponds to an individual animal. Group medians are indicated by vertical bars. **E)** Estimated probability distribution function of θ for superficial and deep SC inputs.

Next, we examined the laminar distribution of SC inputs to each HVA-projecting population (Fig. 4c-e). Here, we find SC inputs in both superficial and deep lamina (Fig. 4b-d). Inputs, especially to AL-, RL-, and AM-projecting pulvinar neurons, were often found near the border of the optic (op) and intermediate gray (ig) layers, similar to a group of projections in squirrels described by Fredes et al. that target the ipsilateral rostral pulvinar (Fredes et al., 2012). Overall, we find a low proportion of inputs in the sg layer compared to previous studies using non-specific retrograde tracers in the pulvinar (Roth et al., 2016; Scholl et al., 2021; N. Zhou et al., 2017). Since the sg layer projects to the caudal pulvinar (or the inferior pulvinar in primates), this discrepancy is also best explained by incomplete spread of the helper AAVs in thalamus. (Abramson & Chalupa, 1988; Beltramo & Scanziani, 2019; Benevento & Standage, 1983; Bennett et al., 2019; Harting et al., 1972; Lyon et al., 2010; Stepniewska et al., 2000; N. Zhou et al., 2017). Nevertheless, our results demonstrate a potential extrageniculate visual pathway to all HVAs in the mouse, which may be less specialized than that of primates (Diamond, 1976; Harting et al., 1973; Petry & Bickford, 2019; Snyder & Diamond, 1968).

Benavidez et al. demonstrated that corticotectal projections from different HVAs are topographically organized along a polar axis in the SC (2021). It is unknown, however, if the SC neurons projecting to different pulvinar→HVA pathways are similarly organized along this axis. To quantify the spatial distribution of these inputs, we registered each brain to a common reference and generated polar coordinates for every rabies-labeled SC neuron. In both superficial and deep layers, inputs to LM- and PM-projecting pulvinar neurons were concentrated in medial SC, whereas inputs to AM-, RL-, and AL-projecting pulvinar neurons were more broadly distributed, extending into the centromedial and centrolateral divisions described in Benavidez et al. (Fig. 4e)(2021). This organization of inputs is consistent with the preferred elevations of the target HVAs, and it also suggests that pulvinar projections to areas AM, RL, and AL may receive multimodal input through the central divisions of the SC.

### The pulvinar relays multimodal information to visual cortex

We next looked at cortical inputs to HVA-projecting pulvinar neurons at the resolution of different cortical areas (Fig. 5). The cerebral cortex is the largest source of input to the pulvinar, and this input originates from a wide range of sensory, association, and frontal regions (Cappe et al., 2009; Froesel et al., 2021; Gattass et al., 1978). Unsurprisingly, we find that the plurality of the input to HVA-projecting pulvinar neurons is visual, followed by retrosplenial (RSP), somatosensory (SS), motor, and auditory (AUD) input (Fig. 5c). When considering the functional significance of these inputs, the distribution of L5CT “driving” cells will reflect the information that is likely to be relayed. We classified cortical inputs into layers based on their locations relative to the boundaries of a L5 marker, Ctip2 (Fig. 5a). Consistent with the reported proportions of these two CT cell types (de Souza et al., 2021), we observed nine times more L6 inputs than L5 inputs. All HVA-projecting groups receive the most L5 input from visual cortex. Interestingly, PM-projecting pulvinar neurons received almost as much L5 driving input from RSP as from visual cortex. RSP is an association area which occupies a higher position than extrastriate cortex in the visual hierarchy. RSP L5 input was substantial for all other cortical targets as well, so this pathway could be an interesting deviation from the hierarchical model depending on the pulvinar cell types involved. Multisensory input to the pulvinar was dependent on cortical target. LM- and PM- projecting neurons received very little input from SS and AUD cortex; and these inputs were relatively high for projections targeting AL, RL, and AM. Motor input to the pulvinar is preferentially routed to medial cortical targets AM and PM.

**Figure 5.**
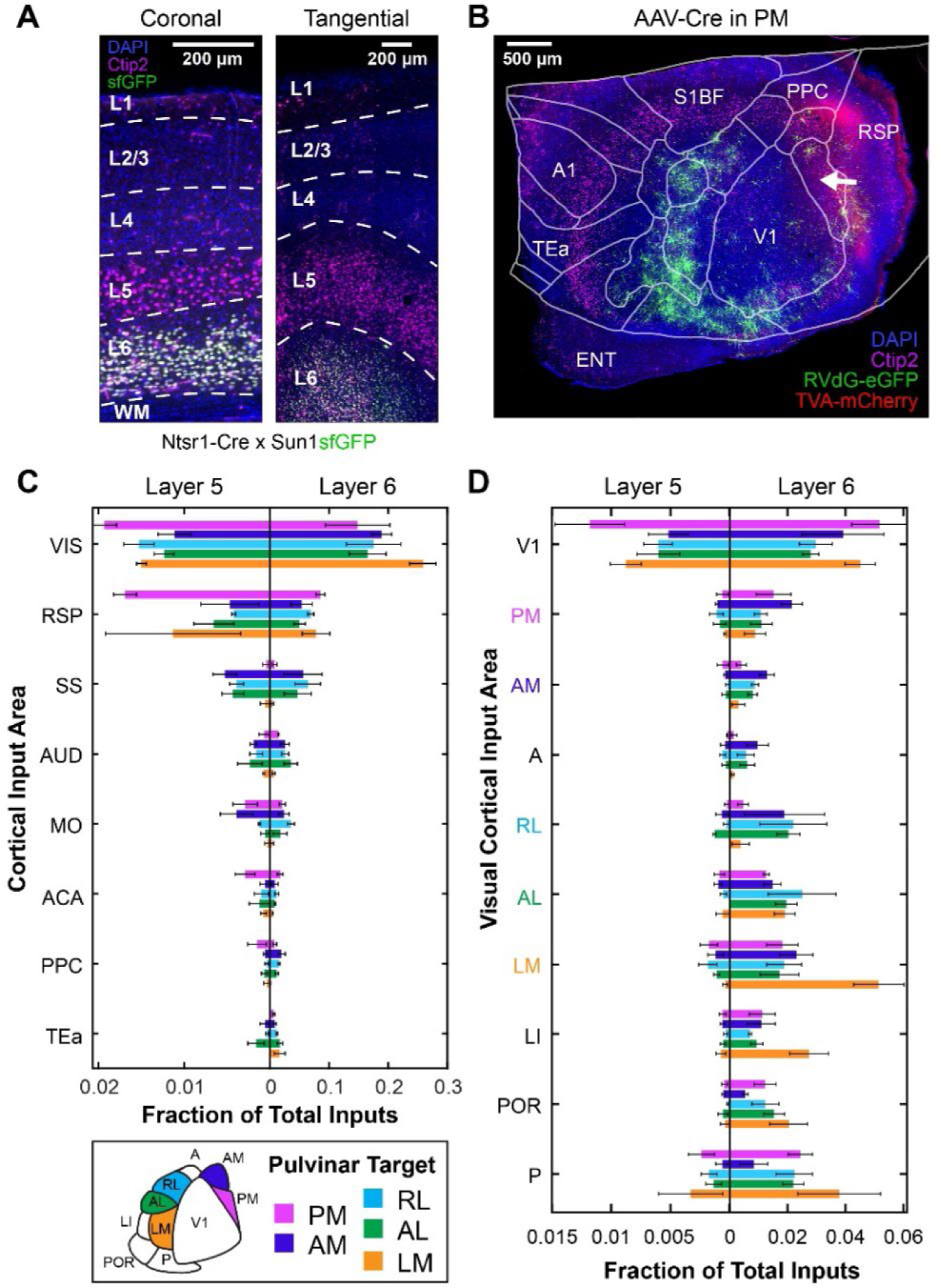
Areal distributions of cortical inputs to pulvino-cortical neurons. A) Immunohistochemical labeling of Ctip2 (magenta) to label L5 in Ntsr1-Cre x Sun1sfGFP transgenic mice expressing GFP in L6CT neurons in coronal (left) and tangential (right) sections. Scale bars = 200 μm. **B)** Example tangential cortical section showing GFP+ inputs to a PM- projecting starter population (animal PM2). Pulvinar axons (red) project to middle layers of medial HVAs. Ctip2 (magenta) defines a band of layer 5. Visual area boundaries from ISI are aligned with cortical sections and merged with the CCF flatmap boundaries of nonvisual areas. Scale bar = 500 μm. **C)** Cortical inputs from layer 5 (left x axis) and layer 6 (right x axis) to five projection populations in the pulvinar expressed as the fraction of total inputs. Bar lengths correspond to the median values from each target area. Error bars show ± median absolute deviations. **D)** Layer 5 (left x axis) and layer 6 (right x axis) inputs from V1 and nine HVAs. Bar lengths correspond to the median values from each target area. Error bars show ± median absolute deviation. ACA: Anterior cingulate area; A1: primary auditory cortex; AUD: auditory cortex; ENT: entorhinal cortex; MO: motor cortex; PPC: posterior parietal cortex; RSP: retrosplenial cortex; S1BF: Primary somatosensory cortex, barrel field; SS: somatosensory cortex; TEa: temporal association areas; V1: primary visual cortex; VIS: visual cortex

### Transthalamic pulvinar CTC connections reflect hierarchical relationships between visual areas

Further separating visual input by area revealed a striking difference in driving CT input from V1 versus HVAs (Fig. 5d). Regardless of the downstream target, L5CT driving input to the pulvinar originates primarily from V1, mirroring the feedforward CC pathways from V1 to HVAs. Shuffled distributions of L5CT and L6CT cells show that independent of target area, HVA inputs to the pulvinar are laterally biased, with increasing input moving from AM→RL→AL→LM and intermediate input from medial area PM (Fig. 7a,b). This trend cannot be explained by HVA size alone, as larger area RL provided relatively little CT input. Consistent with a feedforward CTC relay, LM, which occupies an intermediate level between V1 and other HVAs in the cortical hierarchy (Harris et al., 2019; Wang et al., 2012), provides L5CT input to most other HVA- projecting pulvinar populations (Fig. 5d, Fig. 7c). We also find relatively high proportions of pulvinar input from L5CT neurons in area P, which sends very few direct CC projections to the 5 HVAs targeted here (Wang et al., 2012). Modulating L6CT inputs were more distributed than drivers (Fig. 5d, Fig. 6a), but most cells were still found in V1.

**Figure 6.**
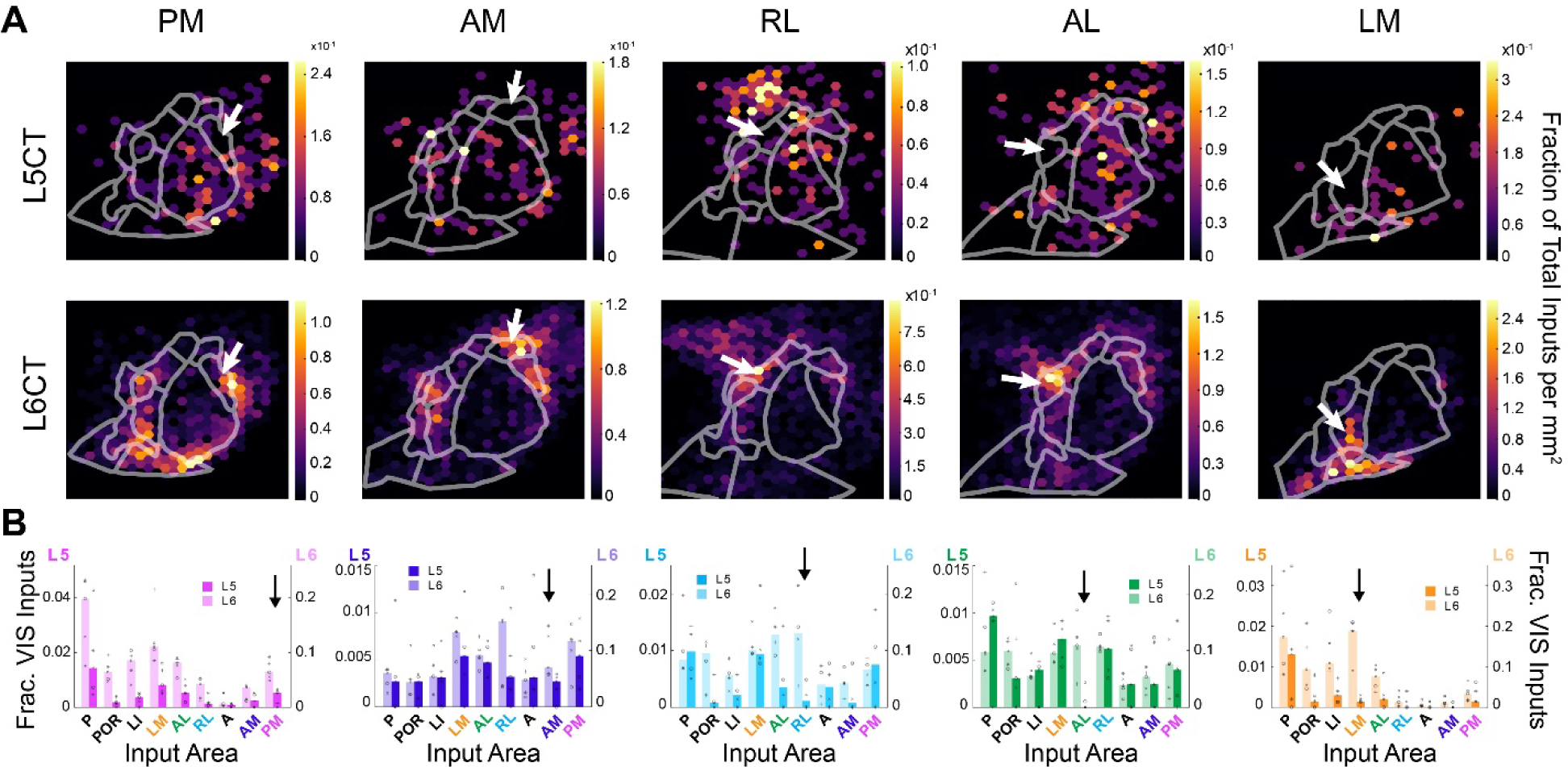
Distribution of driver and modulator input from visual cortex. A) Density of inputs to HVA-projecting pulvinar neurons from L5 (top) and L6 (bottom) for five example animals (from left to right: PM2, AM2, RL2, AL1, LM3). **B)** Higher visual area inputs to each pulvinar projection population from L5 (dark bars, left axis) and L6 (light bars, right axis). Inputs are expressed as a fraction of visual cortical inputs. Arrows denote reciprocal connections.

**Figure 7.**
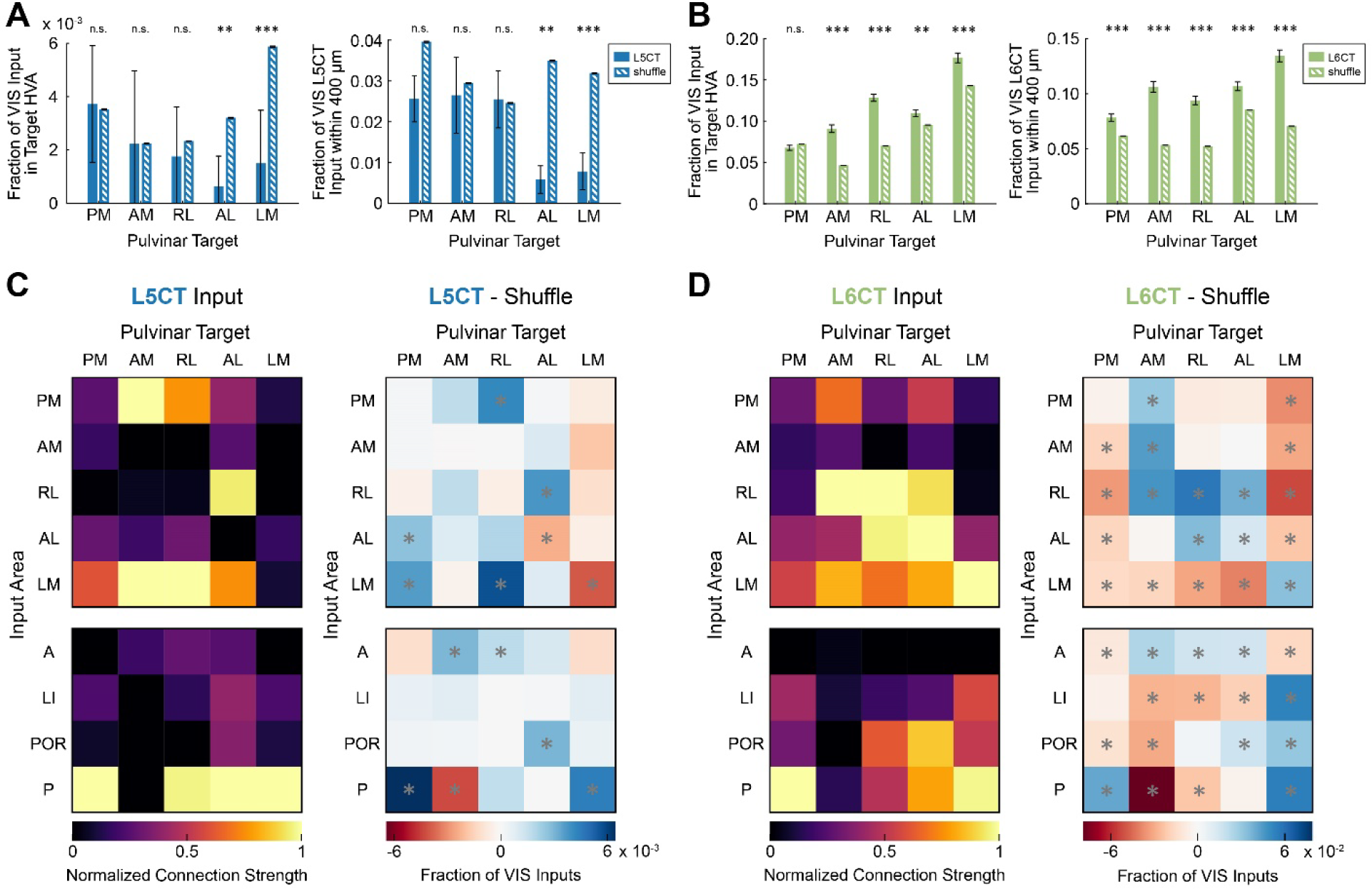
Complementary distributions of driver and modulator CTC pathways. A) Left: Average reciprocal input from L5CT (solid blue bars) to the pulvinar compared to a distribution of L5CT input cells with shuffled targets (hatched blue bars). Input is expressed as a fraction of all visual cortical input. Error bars represent the standard error of the proportion. n.s.: not significant, *: p<0.05, **: p<0.01, ***: p<0.001. Right: Average fraction of visual cortex L5CT inputs within 400 μm of each HVA injection site compared to a shuffled distribution. Error bars represent the standard error of the proportion. **B)** Same as (A) for L6CT input. **C)** Left: Relative strength of HVA L5CT input (rows) to each pulvinar→HVA projection (columns). Median input proportions are normalized to the maximum for each target. The upper matrix shows inputs from traced areas, where reciprocal connections lie along the diagonal. The lower matrix shows additional inputs from HVAs that were not targeted for tracing. Right: Difference between each HVA L5CT connection and the connection expected from a shuffled distribution, expressed as a fraction of all visual cortical inputs. Grey asterisks indicate significant differences (p<0.05 after controlling the false discovery rate of multiple comparisons with the Benjamini-Hochberg procedure). **D)** Same as in (C) for L6CT input.

### HVA input to the pulvinar depends on CT cell type and TC projection target

For input/output relationships among HVAs (Fig. 6b, Fig. 7c-d), both L5CT and L6CT inputs demonstrated significant associations with the target area, indicating that these inputs are not generalized for all projection targets (L5CT: p < 1.0 x 10^-4^, Fisher’s exact test with Monte Carlo simulation; L6CT: χ^2^ = 1.0 x 10^3^, p = 9.6 x 10^-190^, Pearson’s χ^2^ test of independence). Further inspection of projection-specific input distributions revealed complementary organization of L5 and L6 inputs from HVAs. L5 inputs were rarely seen near the target area, but L6 cells were concentrated near the injection site (Fig. 6, Fig. 7b). Except for area PM, reciprocal connections from L6 were overrepresented compared to a null distribution of inputs (PM: p = 0.83, AM: p < 1.0 x 10^-4^, RL: p < 1.0 x 10^-4^, AL: p = 4.1 x 10^-3^; LM p < 1.0 x 10^-4^, permutation test adjusted with Benjamini-Hochberg procedure; Fig. 7b,d, See Methods). L5, however, rarely made reciprocal connections back to its pulvinar projection neurons (Fig. 7a,c 5b). Comparison of L5CT reciprocal connections to the shuffled distribution revealed that AL- and LM- projecting neurons received statistically fewer reciprocal inputs than would be observed by chance (AL: p = 3.7 x 10^-3^, LM: p = 0.5 x 10^-3^, permutation test with Benjamini-Hochberg procedure). PM-, AM-, and RL-projecting reciprocal inputs were not statistically different than the shuffled distribution (PM: p = 0.62, AM: p = 0.62, RL: p = 0.46, permutation test with Benjamini-Hochberg procedure). When present, L5CT reciprocal HVA inputs were typically only one or two neurons, so proportions are highly sensitive to small deviations in HVA borders or alignment.

To investigate the anti-reciprocal L5CT trend with a more robust measure, we analyzed the continuous spatial distributions of inputs. Compared to a distribution of cells with shuffled target areas, L5CT inputs were significantly less likely to be found within 400 μm of the injection site for target areas AL and LM (AL: p = 0.001, LM: p = 4.2 x 10^-3^, permutation test with Benjamini-Hochberg procedure). For target areas PM, AM, and RL, L5CT inputs within 400 μm were not significantly different than the shuffled distribution (PM: p = 0.11, AM: p = 0.53, RL: 0.53, permutation test with Benjamini-Hochberg procedure; see Fig. 7a).

L6CT inputs, conversely, were statistically more likely to be found near the injection site for all target areas (p < 1.0 x 10^-4^, permutation test with Benjamini-Hochberg procedure, see Fig. 7b). L5CT drivers and L6CT modulators therefore form complementary CTC pathways between higher order cortex and thalamus, similar to the CT connections between V1 and dLGN.

## Discussion

Understanding how higher-order thalamic nuclei contribute to cognitive functions like attention, decision-making, and motor planning requires a detailed description of their input/output relationships with the cortex (Halassa & Sherman, 2019). Using projection-specific rabies tracing, this study establishes the first comprehensive map of CTC synaptic connections in the mouse pulvinar. We identify complementary circuit motifs for driving and modulating CT input (Fig. 8). Consistent with two prevailing models of CTC organization, L5CT “driver” inputs form hierarchical, transthalamic relays between V1 and all HVAs, whereas L6CT “modulator” inputs prefer reciprocal CTC loops (Sherman & Guillery, 2011, 2013; Shipp, 2003). We also demonstrate a disynaptic pathway from the SC to *all* HVAs in the mouse.

**Figure 8.**
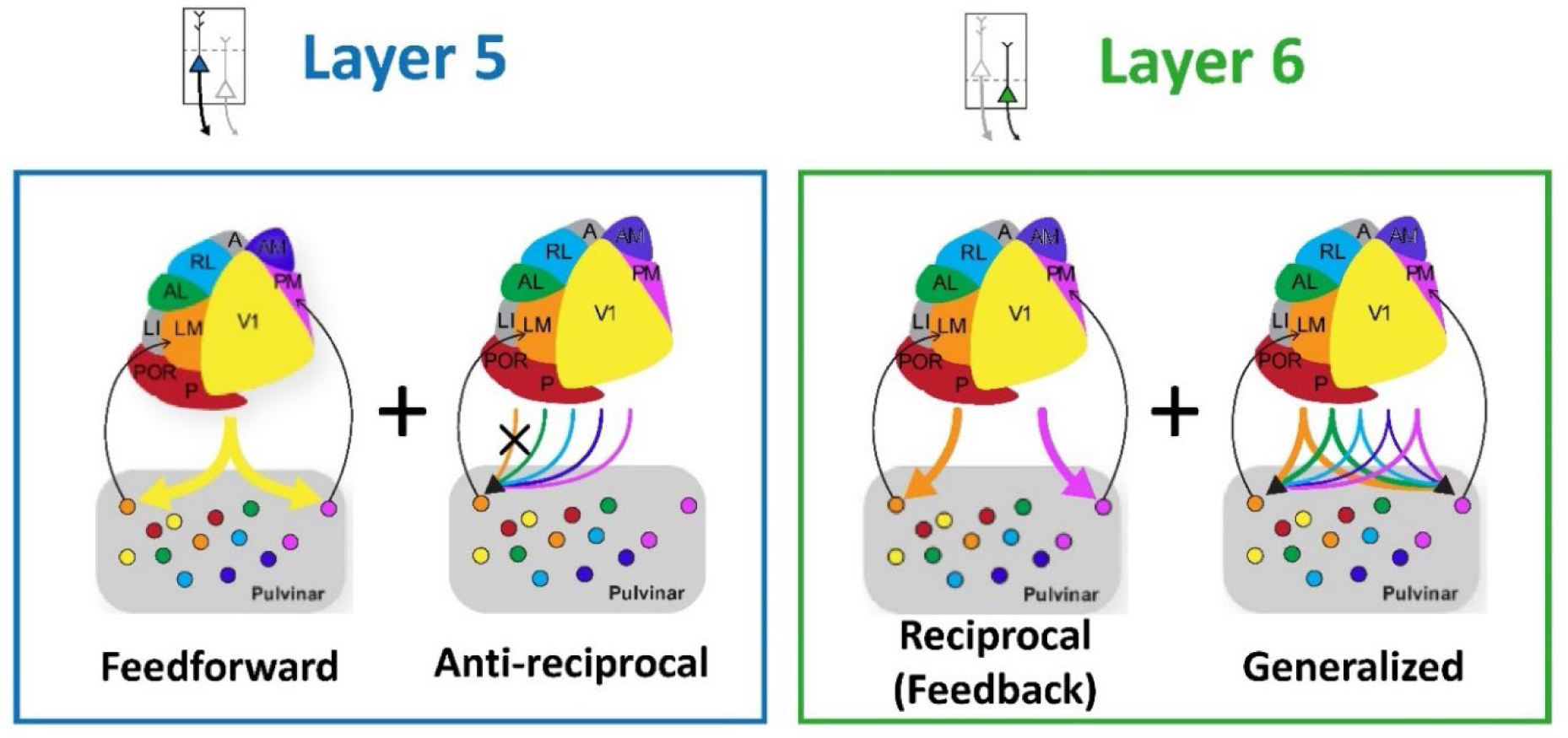
Summary of Input/Output Connections in the Pulvinar. Left: Schematic showing connections of L5CT projections with the pulvinar. Inputs from L5CT drivers are feedforward and anti-reciprocal. Right: Schematic showing L6CT connections with the pulvinar. L6CT modulators are distributed and reciprocally-biased.

CTC pathways can act as parallel routes for signal transmission between connected cortical areas (Guillery, 1995, 2003, 2005; Sherman & Guillery, 2011, 2013). Since only L5CT projections are capable of driving strong postsynaptic responses in pulvinar neurons, such transthalamic relays are restricted to CTC pathways originating from L5 of upstream cortical areas. Several recent studies have confirmed driving connections from V1 to AL- and PM-projecting pulvinar neurons (Blot et al., 2021) and from the visual cortex to ACA-projecting pulvinar neurons (Leow et al., 2022). Similar higher-order transthalamic circuits exist from S1 to S2- and M1-projecting POm neurons (Mo & Sherman, 2019; Theyel et al., 2010). These individual connections suggest that feedforward, transthalamic relays are a general organizational feature of cortical networks, but until now there has not been a comprehensive map input/output of CTC connections for an entire cortical hierarchy. We found that L5CT inputs to all pulvinar→HVA projections originate predominantly from V1, confirming that parallel, transthalamic relays are a general organizational principle. Leow et al. extend this framework beyond sensory cortex with input/output tracing of pulvinar→ACA projections, where they find more L5 inputs from HVAs than from V1 (Leow et al., 2022). This suggests that L5CT distributions reflect true hierarchical relationships rather than trivial factors such as area size. Additional input/output tracing of pulvinar projections to areas TEa, RSP, and other higher-order sensory cortices could reveal additional pathways from HVA L5CT cells.

Compared to V1, HVA L5CT inputs were rare. Nevertheless, comparison of L5CT inputs *between* HVAs revealed a significant absence of reciprocal connections when compared to a shuffled distribution for AL- and LM- projecting cells. While the lack of reciprocal connections to lateral HVAs is unlikely to have a strong influence on the overall activity of TC projections, the wiring specificity is an interesting developmental phenomenon and likely involves active signaling mechanisms (Grant et al., 2012). This finding provides direct support for the “no-strong-loops” hypothesis, which asserts that there should be no reciprocal connections with strong synaptic weights in both directions (Crick & Koch, 1998). While the L5CT projections are “strong”, we do not yet have a method to experimentally separate the two classes of TC neurons in the mouse. The pulvinar contains a mix of core and matrix cells (Jones, 1998; Nakamura et al., 2015; Rockland, 2019; Rockland et al., 1999), and only the former would constitute a “strong” projection.

Pulvinar→HVA projections are predominantly core neurons (Marion et al., 2013; Rockland et al., 1999; Rockland, 2019; N. Zhou et al., 2018), but the ratio of core and matrix projections varies across pulvinar subdivision (Nakamura et al., 2015). The few reciprocal L5CT connections that we do find for areas RL, AM, and PM could be inputs to the matrix-enriched rostromedial pulvinar, which would not violate the “no-strong-loops” hypothesis, as the CTC loop would terminate in a weak projection. Indeed, Miller-Hansen and Sherman identified significant reciprocal L5CT inputs to pulvinar→V1 projections, which are exclusively matrix cells (2022). In nuclei like the pulvinar where TC cell types are mixed, understanding the differential organization of core and matrix CTC circuits in nuclei like the pulvinar where they are mixed will require the identification of genetic markers.

Previous input tracing of the pulvinar→HVA projections did not find a significant lack of reciprocal connections from L5CT cells in AL (Blot et al., 2021). We suggest several technical factors related to viral targeting and HVA border measurement that could explain this discrepancy. In our control experiments with no AAV-DIO-oG, we found that high concentrations of AAV- FLEx-TVAmCherry caused significant retrograde infection of L5CT neurons in the target area (Fig. 2). Without such control experiments, we cannot rule out the possibility that previously observed, reciprocal L5CT labeling was a false positive artifact from direct rabies uptake by CT axon terminals with Cre-mediated expression of TVA. Since the overall number of L5CT inputs is low, this target-specific artifact could obscure the true anti-reciprocal distribution. Additionally, large injection volumes of AAV-Cre that spread across areas can result in labeling of L5CT inputs from adjacent HVAs that might target the other, as we observed for injections which crossed the P/POR border (data excluded, see methods). Thirdly, HVA borders vary substantially between animals, so registering cells to a common atlas will inevitably lead to some assignment errors, which are more detrimental for the L5CT population, which often has a low total cell count.

Consistent with bulk tracing studies, L6CT cells targeting specific pulvinar→HVA projections comprise almost 90% of total cortical input, and their distribution follows the general topography of all L6CT inputs to the pulvinar (Bennett et al., 2019). As expected from the overlap of cortical input/output regions in the pulvinar, we find a bias for reciprocal L6CT connections. Although reciprocal loops are more likely than chance, they are rarely the largest L6CT input to a given pulvinar→HVA projection. Regardless of target HVA, the L6CT pathway from V1 is predominant.

While we could not delineate L6 sublaminae in our tangential preparation, previous work has shown that V1 L6CT input to the pulvinar arises from lower L6a and L6b, whereas L6CT input from HVAs also includes upper L6a cells (Bourassa & Deschênes, 1995; Roth et al., 2016). In the somatosensory system, these L6CT subtypes form functionally distinct circuits. (Frandolig et al., 2019; Whilden et al., 2021). Further splitting HVA L6CT projections by sublayer, then, might reveal additional differences in circuit organization and function. L6CT modulator inputs do not meaningfully influence the receptive field content of thalamic neurons, but they are capable of inhibiting thalamic activity through their collaterals to the TRN and of controlling the gain and frequency of thalamic sensory responses (Bourassa & Deschênes, 1995; Crandall et al., 2015; Cruikshank et al., 2010; Kirchgessner et al., 2020, 2021). Most of these effects have been described for L6CT feedback to first-order thalamic nuclei. Evidence shows that these modulatory influences are similar for the pulvinar, but our results demonstrate that the sources of L6CT modulation are much more diverse than for corticogeniculate feedback. Broad L6CT modulation to all pulvinar projections could be one mechanism supporting the synchronization of cortical areas by the pulvinar (Contreras et al., 1996; Saalmann et al., 2012; H. Zhou et al., 2016).

Most retinal input reaches the cortex via the dLGN; however, the pulvinar provides a secondary pathway for incoming retinal signals via the SC (Baldwin et al., 2011; Benevento & Standage, 1983; Berman & Wurtz, 2008; Fish & Chalupa, 1979; Lyon et al., 2010; Petry & Bickford, 2019). This circuit is important for cortical development (Bourne & Morrone, 2017; Homman-Ludiye & Bourne, 2019), and it can support “blindsight”, a phenomenon whereby visually-guided actions remain intact even after lesions of striate cortex cause perceptual blindness (Bertini et al., 2013, 2018; Kinoshita et al., 2019; Pöppel et al., 1973; Sanders et al., 1974; Stoerig, 1997; Takakuwa et al., 2021; Weiskrantz et al., 1974). In the macaque, this pathway only targets dorsal visual areas (Lyon et al., 2010). In a notable species distinction, we found that all HVAs in the mouse receive disynaptic input from the SC, which originated from both superficial and deep layers. While the framework of dorsal and ventral pathways has been highly influential in the study of primate visual cortex (Goodale & Milner, 1992; Kaas & Baldwin, 2019; Kaas & Lyon, 2007; Mishkin et al., 1983; Ungerleider, 1982), the classification of two distinct visual streams in the mouse is less clear (Beltramo & Scanziani, 2019; Berman & Wurtz, 2008; Kaas & Baldwin, 2019; Wang et al., 2011, 2012). Our results suggest that tectal input may be more relevant to the mouse cortex than previously thought, and that all mouse HVAs have some degree of connectivity resembling the primate dorsal pathway from SC.

In summary, we have generated a comprehensive input/output map of the mouse pulvinar that extends the principles of first order corticothalamic connectivity to higher-order nuclei. Our data confirm long-standing theories that driving CT inputs are feedforward relays and avoid reciprocal loops (Crick & Koch, 1998; Sherman & Guillery, 2011), whereas L6CT inputs are distributed and reciprocally-biased (Fig. 8). These findings support the distinction of direct and transthalamic pathways between cortical areas, but additional research is necessary to investigate the functional roles for these circuits.

## Methods

### Animals

Thirty-three Adult female and male C57BL/6J mice (The Jackson Laboratories) aged 6-10 weeks were used in this study. All experimental procedures followed procedures approved by the Salk Institute Animal Care and Use Committee.

### Viruses

AAV5-Ef1α-Cre-WPRE (2.22 x 10^12^ GC/mL; UNC Vector Core) AAV8-CAG-FLEx-TCB (2.04 x10^13^ GC/mL; Salk GT3 Core) AAV8-CAG-FLEx-oG-WPRE-SV40pA (1.55 x10^13^ GC/mL; Vigene Biosciences) EnvA+RVΔG-eGFP (3.29 x10^8^ TU/mL; Salk GT3 Core) To trace the inputs to specific projection populations, a mixture of AAV8-CAG-FLEx- TCB (AAV-FLEx-TVA) and AAV8-CAG-FLEx-oG-WPRE-SV40pA (AAV-FLEx-oG) was injected into the pulvinar. To prevent glycoprotein-independent retrograde labeling of CT neurons, the AAV-FLEx-TVA was diluted to between 8.16x10^11^ - 4.04x10^12^ GC/mL with Hank’s Buffered Saline Solution, then mixed in a 1:1 ratio with undiluted AAV-FLEx-oG. In control experiments with no AAV-FLEx-oG, higher concentrations of AAV-FLEx-TVA resulted in non-specific RVΔG-eGFP expression in L5b and L6 of cortex, restricted to the location of the AAV5-Cre injection site (Fig. 2). At the optimized TVA concentration, RVΔG-eGFP+ neurons were observed locally at the pulvinar injection site, but not in the cortex or any other distal structures. We frequently saw a small number of GFP+ astrocytes at the AAV-Cre injection site, usually in upper cortical layers and with distinct morphology. These were present regardless of AAV-FLEx-TVA concentration and were not counted in the experimental brains. Control injections without Cre but including glycoprotein did not result in any long-range labeling, even at the higher TVA concentrations which failed the no-glycoprotein control.

### Surgical Procedures

Mice were anesthetized with isoflurane (2% induction; 1.5% maintenance) and mounted in a stereotax (David Kopf Instruments Model 940 series). Using a carbide burr, a small craniotomy was made over the left pulvinar (-1.8 to -2.3 from bregma; -1.2 to -1.6 lateral from the midline). Injection coordinates varied depending on the cortical projection region. The virus was pressure-injected via syringe at a rate of 10 nL/min through a tapered glass pipette (25-30 μm tip inner diameter) at a depth of 2.3-2.5 mm below the pial surface. To prevent backflow of the virus into the hippocampus and cortex, the pipette was left in place for at least 10 minutes before retraction. During the same surgery, a custom metal headframe was attached to the skull with dental cement (C&B Metabond, Parkell) as previously described (Juavinett et al., 2017). Either buprenorphine-SR (0.5-1 mg/kg) or meloxicam (4 mg.kg) were administered subcutaneously for post-op analgesia. Mice recovered for 1-4 days in their home cage with a suspension of ibuprofen (30 mg/kg) in water prior to intrinsic signal imaging (ISI).

After ISI mapping, blood vessel landmarks were used to target injections of AAV5-Ef1α-Cre-WPRE to one of six HVAs (PM, AM, RL, AL, LM, or POR). We chose AAV5 for its high propensity to infect long-range thalamocortical axons. In comparison, AAVretro-Cre caused marked toxicity and reduced cell labeling at the cortical injection site, and when diluted to nontoxic concentrations, its efficient was comparable to AAV5-Cre. The virus was at two depths between 200-500 μm below the pial surface. The total volume of AAV-Cre ranged between 35-75 nL and was chosen based on the size of each HVA to maximize coverage without spreading into adjacent visual areas. The skull thinning required for ISI is appropriate for short survival times but leads to infection and tissue degradation after approximately a week. Due to the long survival time required for expression of oG, we removed the skull after thinning and placed a molded cement cover over the craniotomy to preserve the tissue health. A subset of animals was imaged without any skull thinning to guide injections, and then subsequently re-imaged with skull thinning prior to euthanasia.

Three weeks after injection of AAV-Cre, 350 nL of EnvA+RVΔG-eGFP was injected into the pulvinar at the previous site at 2-3 depths between 2.3-2.6 mm below the pial surface. Buprenorphine-SR (0.5-1 mg/kg) was administered subcutaneously, and mice received ibuprofen in their water (30 mg/kg) as they recovered in their cages for 10 days until tissue harvesting.

### Intrinsic Signal Imaging

After viral injections in the thalamus, higher visual areas (HVAs) in cortex were identified using intrinsic signal imaging (ISI) through a thinned skull as previously described (Garrett et al., 2014; Juavinett et al., 2017). Mice were anesthetized with 0.1-1% isoflurane and sedated with (1 mg/kg) chlorprothixene hydrochloride injected intramuscularly. Anesthesia was continuously monitored and adjusted to maintain a lightly anesthetized state (1-1.5 breaths/second). Most experiments resulted in maps with seven visual areas (V1, PM, AM, RL, AL, LM, and P/POR) successfully segmented. These maps were used to guide cortical injections and to align post-mortem tissue to the HVA borders. Occasionally, the initial map of visual cortex identified a subset of these areas. In these cases, the partial map was sufficient to target a cortical injection, and ISI was repeated just prior to the experimental endpoint to obtain a complete map that was suitable for post-mortem tissue alignment. The border between P and POR was usually ambiguous, so these injections were targeted generally to the positive field sign area directly behind LM, which likely included both areas.

### Exclusion Criteria

Since the pulvinar is topographically organized with respect to its cortical inputs and outputs, the accuracy of targeting for both helper virus injection and the RVΔG injection was crucial to avoid biasing the measured input samples. Animals were excluded if the two injections did not overlap, in which cases we observed reduced starter cell populations and little to no transsynaptic labeling. We excluded animals in which starter cell locations were restricted or heavily biased to the tectal- recipient caudal Pulvinar, as the corresponding cortical inputs were either absent or only present in caudally-projecting area POR. We also excluded animals in which starter cells were biased compared to the overall distribution of starter cells for a given projection population, which aligned with the topographical organization that has been previously reported (Bennett et al., 2019; Juavinett et al., 2020). We excluded rare experiments in which surface blood vessels could not be accurately aligned. While we injected at least 6 mice that met these criteria for areas P and POR, our injections were not restricted to the individual areas and instead covered both P and POR. Due to this ambiguity in cortical target, these experiments were also excluded from analysis.

### Histology

Mice were euthanized with an intraperitoneal injection of Euthasol (15.6mg/ml), then transcardially perfused with phosphate-buffered saline (PBS) followed by 4% paraformaldehyde (PFA). After dissection, brains were post-fixed in 2% PFA and 15% sucrose in PBS for 16-24 hours at 4°C, then submerged in 30% sucrose for an additional 16-24 hours at 4°C.

The left visual cortex and hippocampus were dissected from the rest of the brain and sectioned tangentially on a freezing microtome. After the first 250 μm section which contained the surface blood vessels, the remaining cortical tissue was sectioned in 50 μm increments. Immunohistochemical staining of the free-floating cortical sections amplified the eGFP and mCherry signals and provided a laminar marker (Ctip2) to distinguish L5 from L6. After blocking at room temperature in PBS with 10% normal donkey serum (NDS) and 0.5% Triton-X, tissue was incubated for 16 hours at 4°C with goat anti-GFP (1:500; 600-101-215; Rockland Immunochemical), rabbit anti-dsRed (1:500; 632496; Takara Bio USA), and rat anti-Ctip2 (1:1000; ab18465; Abcam) in PBS with 1% NDS and 0.5% Triton-X. The tissue was then incubated for 6 hours at room temperature with donkey anti-goat conjugated to Alexa Fluor 488 (1:500; A-11055; Thermo-Fisher), donkey anti-rabbit conjugated to Alexa Fluor 568 (1:500; A- 10042; Thermo-Fisher), and donkey anti-rat conjugated to Cy5 (1:500; 712-175-153, Jackson ImmunoResearch) in PBS with 1% NDS and 0.5% Triton-X, followed by a counterstain with 10 μM DAPI in PBS. Tangential sections were mounted on gelatin-subbed slides, air-dried, then coverslipped with polyvinyl alcohol mounting medium containing 1,4-diazabicyclo-octane (PVA-DABCO).

The remaining brain tissue was sectioned coronally on a freezing microtome in 50 μm increments from the olfactory bulb to the end of the cerebellum. Free-floating sections were blocked in PBS with 5% NDS and 1% Triton-X for 1 hour at room temperature, then incubated with goat anti-GFP (1:500; 600-101-215; Rockland Immunochemical) and rabbit anti-dsRed (1:500; 632496; Takara Bio USA) in PBS with 1% NDS and 0.5% Triton-X for 16-24 hours at 4°C. The tissue was then incubated in the appropriate secondary antibodies for 2 hours at room temperature, then counterstained with DAPI. Coronal sections were mounted on glass slides with PVA-DABCO.

### Image Processing and Registration

All slides were imaged on an Olympus BX63 epifluorescence microscope with a 10X objective. Cells in coronal sections were counted manually and assigned to brain areas based on cytoarchitectonic landmarks by an expert, or automatically registered to the Allen Common Coordinate Framework (CCF) (Wang et al., 2020) after transforming sections using the SHARP-TRACK program (Shamash et al., 2018) and custom Matlab scripts. Medial TRN cells were sometimes mistakenly assigned to VPL due to slight misalignments at the nuclei border. Therefore, those cells were manually checked and compared to cytoarchitectonic landmarks. The percentage of cells assigned to major brain areas was comparable between these two registration methods.

Z-stacks of tangential cortical sections were processed in the Olympus cellSens software using the Extended Focal Imaging method, then converted for additional processing using ImageJ software (NIH). Cortical sections were virtually “flattened” using radial blood vessel alignment (Kim et al., 2020), which corrected for the curvature of the cortex and aligned cells in deep layers with the HVA assignments defined at the pial surface. This alignment was achieved with sequential affine matrix transformations of deeper sections onto superficial sections using the Landmarks Registration plugin. When apical dendrites could be matched between images, they were included as landmarks in addition to radial blood vessels. The anatomical reference and HVA border map obtained during ISI were similarly warped onto the aligned image stack using common surface blood vessel landmarks present in the top tangential section. Cells in each transformed section were registered to the aligned HVA map and assigned to a cortical layer based on the borders of the Ctip2 signal in individual sections. The retinotopically organized, positive field sign patch designated as RL usually extended into the S1 barrel field as has been previously reported (Zhuang et al., 2017), so the anterior border was modified from the automatically segmented borders so that it aligned with the posterior edge of the imaged S1 barrel fields. All RL injection sites were still contained within the truncated borders. Area A lies in between RL and AM (Garrett et al., 2014; Wang & Burkhalter, 2007), but was never segmented in our ISI. The medial border of RL was therefore estimated to lie along a line extending from the gamma barrel in between rows C and D to V1. Although the definition of area A and AM relative to the posterior parietal cortex (PPC) is still currently under debate (Gilissen et al., 2021; Lyamzin & Benucci, 2019), we defined A as the region between RL and AM which is visually responsive and retinotopically organized. The area outside of these functional borders and posterior to somatosensory cortex was defined as PPC.

Input cells were frequently labeled in the nonvisual areas contained in tangential sections. To assign these areas, we used a 3D volume created from the registered coronal sections to define the edges of the dissected cortex in the Common Coordinate Framework (CCF). This dissection boundary was then mapped onto the CCF flatmap, which was aligned to the cut edges of the tangential stack.

Visual areas P and POR were only partially mapped by intrinsic signal imaging due to these areas extending behind the lambda suture and, in the case of POR, laterally out of the imaging ROI. The anterior borders with LM and V1 were confirmed with ISI mapping, but the remaining boundaries of POR were defined by the transformed flatmap and the border of the entorhinal cortex which was evident from the cytoarchitecture. We observed a positive field sign band extending behind V1 as has been reported previously (Zhuang et al., 2017). In post-mortem tissue, this region frequently had TVAmCherry^+^ pulvinar axons in layer 4, which confirms that this area is separate from V1, which only receives matrix projections to layer 1 and 5a (N. Zhou et al., 2018). Rather than defining the whole posterior edge of the cortex as area P, we used the most lateral edge of the dense DAPI band in RSPv layer 2/3 to delineate the border between P and RSP.

### Analysis and Statistical Methods

Because the glycoprotein was not labeled with a fluorescent marker, we considered any GFP+ cell near the injection site to be a putative starter cell. All cells outside of this region were classified as inputs and expressed as a fraction of the total input population for each brain region (total inputs summarized in Table 1). Brain-wide inputs to each projection population were compared with a Kruskal-Wallis test, with post-hoc Dunn-Šidák correction for significant comparisons.

To visualize the relative positions of SC inputs to different projection populations (Fig. 4c), cells were registered to the CCF using the forward transforms obtained from tissue registration described previously. For each target HVA, we pooled cell positions across animals without weighting so that brains with less efficient SC input labeling did not skew the distribution. These cells were divided into four 500 µm bins covering the anterior/posterior extent of SC and plotted with the atlas boundaries obtained at the center of each bin (-3.05, -3.55, -4.05, -4.55 mm from bregma).

Benavidez et al. described a systematic organization of SC inputs and outputs that varies along the radial extent of SC (Benavidez et al., 2021). To determine whether the SC inputs to pulvinar share this trend for cortical topography, we defined an angle, theta (θ), representing cell positions along a polar axis where the ventral boundary of the SC bin is 0° and the midline is 90°. Probability density functions were fit to the theta values for the pooled, binned samples of SC cells using a kernel smoothing function with boundary correction at 0° and 90°.

L5CT and L6CT density maps were generated by binning the continuous distributions of cell positions for each animal in hexagonal bins with a width of 180.6 μm. Example brains for each target area are shown. Cartesian distances of input cells to the injected area were calculated from the transformed tissue.

A null distribution of cell distances was generated by shuffling the target area 10,000 times and recalculating the cell distances from the shuffled target locations. To test whether observations of L5CT or L6CT cells near the injection site were different than would be observed by chance, we compared the true proportion of cells within a 400 μm radius to the proportions calculated from the shuffled distributions. We calculated one-tailed p-values for L5CT and L6CT to test for anti- reciprocity and reciprocity, respectively. P-values were adjusted using the Benjamini-Hochberg procedure for control of false discovery rate. Any p-values calculated as 0.0000 are reported as p < 1.0 x 10^-4^.

Outliers with large proportions of inputs in SC conversely had lower overall proportions of visual cortical input, so we normalized HVA inputs to the total visual cortical inputs for discretized area comparisons. Fractions of visual cortical input to each projection target were compared for L5CT and L6CT. To visualize the relative strengths of these HVA connections, we plotted the median input values in a heatmap. Color scales normalized for each target area (column), resulting in a visualization of the relative strengths of each input compared to the strongest input, in arbitrary units.

To test for significant input/output associations for L5CT connections, we performed a fisher’s exact test on the raw cell counts with monte carlo simulation (9.6x10^7^ shuffles) to a estimate a p- value ±0.0001 with 95% confidence. L6CT sample sizes were much higher than L5CT, so a Pearson’s χ^2^ test for independence was applied to the L6CT input/output matrix. The shuffling procedure for cell distances was repeated for the cells’ HVA assignments to generate a null distribution of area counts to compare against projection-specific distributions.

